# Micro topographical instruction of bacterial attachment, biofilm formation and *in vivo* host response

**DOI:** 10.1101/2020.10.10.328146

**Authors:** Manuel Romero, Jeni Luckett, Grazziela P. Figueredo, Alessandro M. Carabelli, David Scurr, Andrew L. Hook, Jean-Frédéric Dubern, Elizabeth Ison, Lisa Kammerling, Ana C. da Silva, Xuan Xue, Chester Blackburn, Aurélie Carlier, Aliaksei Vasilevich, Phani Sudarsanam, Steven Vermeulen, David Winkler, Amir M Ghaemmaghami, Jan de Boer, Paul Williams, Morgan R Alexander

## Abstract

Bio-instructive materials that prevent bacterial biofilm formation and drive an appropriate host immune response have the potential to significantly reduce the burden of medical device-associated infections. Since bacterial surface attachment is known to be sensitive to surface topography, we experimentally survey 2,176 combinatorially generated shapes using an unbiased high-throughput micro topographical screen on polystyrene. This identifies topographies that reduce colonization *in vitro* by up to 15-fold compared with a flat surface for both motile and non-motile bacterial pathogens. Equivalent reductions are achieved on polyurethane, a polymer commonly used in medical devices. Using machine learning methods, a set of design rules based on generalisable descriptors is established for predicting bacteria-resistant micro topographies. In a murine foreign body infection model, anti-attachment topographies are shown to be refractory to *P. aeruginosa* and to recruit a productive host response, highlighting the potential of simple topographical patterning of non-eluting implants for preventing medical device associated infections.

## Body Text

Infections caused by formation of bacterial biofilms on the surfaces of implanted medical devices lead to increased patient morbidity and mortality and constitute a global challenge for the healthcare sector. Indwelling polymer catheters are responsible for the most common healthcare acquired infections, prompting calls from clinicians for technical advances in materials.^1,2^ Additionally, to counteract the rise of multi-antibiotic resistant pathogens, the formation of refractory biofilms on surfaces should be avoided before infections initiate rather than relying on systemic delivery of antibiotics to prevent or resolve infections.

Although once the first choice for patients vulnerable to infection, use of eluting components such as silver ions in implanted medical devices has declined following systematic review and meta-analysis indicating a lack of clinical effectiveness.^3^ The development of non-leachable bio-instructive materials that instruct bacterial and host cells proximal to devices in-service is a promising alternative strategy to reduce infection and improve device acceptance. However, the relationships between cellular responses to materials and their chemical properties are poorly understood, preventing reliable prediction and design of improved materials. Consequently, large diversity screening approaches to identify cell-instructive polymers for a range of biological applications.^4^ Successful approaches include identification of polymers for stimulating stem cell proliferation and phenotype control in regenerative medicine,^5,6^ immune cell instruction for implant device control^7^ and microbial biofilm controlling polymers.^8,9^ Strongly predictive relationships linking molecular descriptors to biological responses have subsequently been identified, allowing the design of polymer chemistries with improved resistance to biofilm formation.^10,11^

In addition to materials chemistry, topography can strongly influence cellular responses to surfaces, with nano and micro topographic cues being successfully used to control human stem cells,^12^ and immune cells,^13^ marine biofouling,^14^ and bacterial surface colonization.^15^ Despite many studies, the lack of a general description of topography and its influence on cellular responses hinders the rational design of textured biomaterials.^16^ Consequently, to identify design rules for topographical control of bacterial responses to surfaces beyond simple geometries or isolated bioinspired topographical designs, we applied an unbiased screen to investigate combinatorially generated shapes using an unbiased high-throughput micro topographical polymer chip screen that has previously been applied to human mesenchymal stromal cells.^17^

### The library of micro topographies

To sample a large range of topographical designs, topographies were comprised of primitives (circles, triangles and rectangles) combined in an algorithm-based approach, which were arrayed to form 300 µm square Topounits separated from each other by 40 µm high walls (**Figs S1 and S2)**.^17^ Two replicates of 2,176 different features were thermally embossed from a silicon master as 10 µm high shapes on each polystyrene TopoChip (**Fig S1**).

### Micro topographical control of bacterial attachment

The screening methodology for measuring the attachment of representative Gram-negative (*Pseudomonas_aeruginosa*) and Gram positive (*Staphylococcus aureus)* bacterial pathogens to the different Topounits is illustrated in **Fig 1a**. A 4 h incubation time and static conditions at 37°C was found to provide a sufficiently stringent assay for the identification of topographies with low and high initial bacterial cell attachment (**Fig S3**). *P. aeruginosa* attachment to each Topounit on the polystyrene TopoChip (n = 22) revealed a wide range of bacterial attachment to micro topographies, plotted in rank order in **Fig 1b**. For *S. aureus*, a similarly broad range of attachment was indicated by the fluorescence intensity differences observed using a separate rank order (**Fig 1b**). Comparison of the response to the two pathogens for all Topounits revealed a surprisingly good correlation, shown in **Fig 1c**, suggesting a common attachment response between two quite different bacterial species, one motile (*P. aeruginosa*) and the other non-motile (*S. aureus*). Interestingly, most micro-patterns on the polystyrene TopoChip showed reduced bacterial attachment with respect to the flat control, especially for *S. aureus* where only ∼4% of the topographies exhibited increased cell attachment compared to flat (**Fig 1b**). These findings suggested that by modifying surfaces with appropriate micro topographies, early cell attachment could be prevented and hence biofilm formation after longer incubation times.

**Figure 1.**
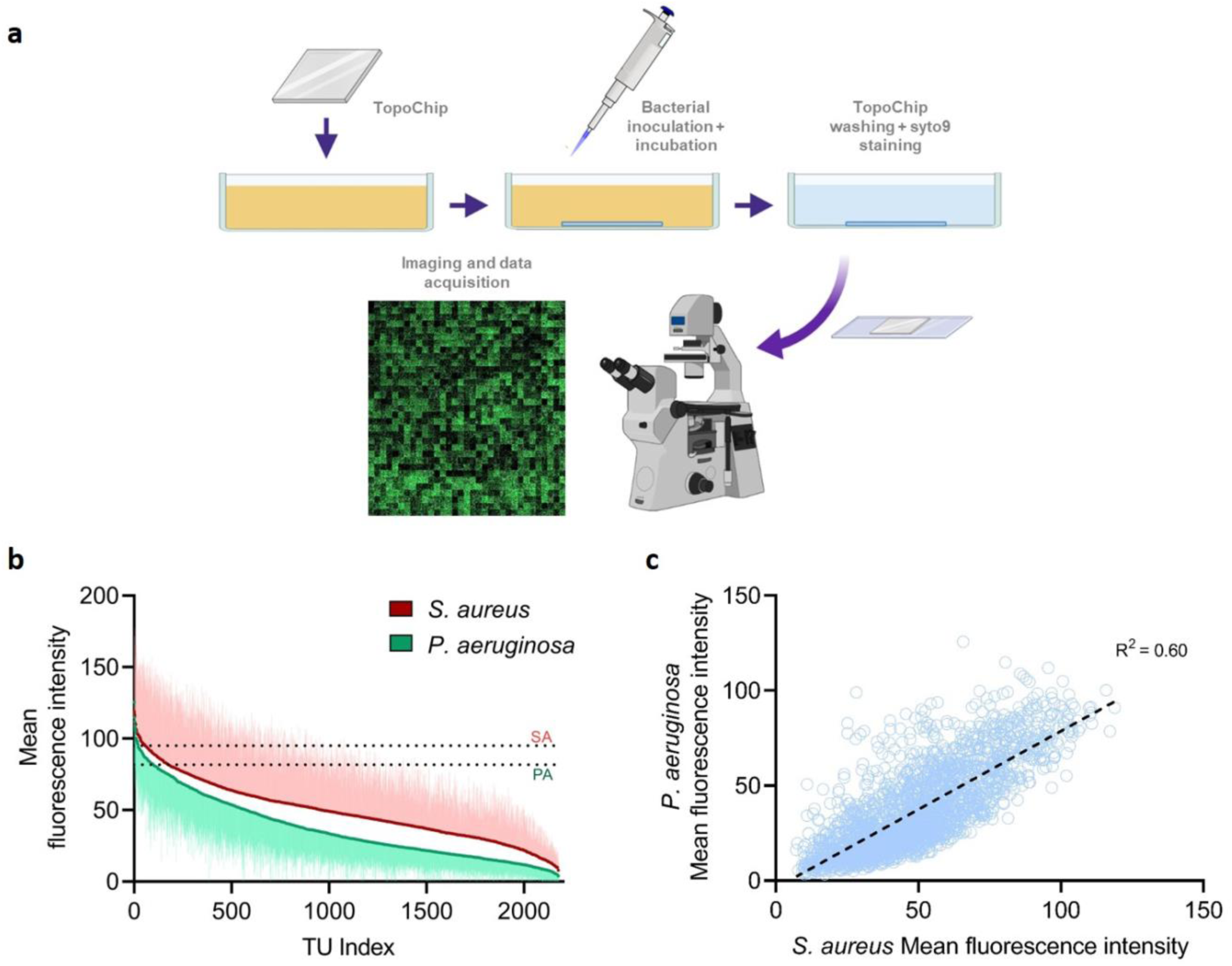
(**a**) Bacterial attachment assay procedure. (**b**) Ranking of topographies according to the mean fluorescence intensity per Topounit after *P. aeruginosa* and *S. aureus* attachment to polystyrene TopoChips after 4-h incubation (n = 22 and n = 6 respectively). The error bars represent one standard deviation. The dotted line corresponds to the mean fluorescence value of the flat surface control for each species. (**c**) mean fluorescence intensities from *P. aeruginosa* versus *S. aureus* for each polystyrene Topounit.

By visual inspection of the high and low-adhering topographies, it appeared that fewer bacterial cells attached to topographies with small gaps between the Topo features (see examples of anti- and pro-attachment Topounits in **Fig S2a**). This was unexpected since the presence of narrow grooves and cavities has been shown to support bacterial retention on surfaces under static growth conditions.^18,19^ To elucidate the critical elements of the Topounit shapes from this rich library, we turned to machine learning to extract generalisable design rules.

### Micro topographical design rules for bacterial attachment

Mathematical descriptors for the topographical shapes were created by co-opting the cell shape analysis functions of the widely used CellProfiler and ImageJ software packages, 242 shape descriptors were obtained from the analysis of the Topounit designs.^13^ Those with Pearson correlations above 0.85 were removed to eliminate co-variant descriptors, resulting in 68 that were used to train models of *P. aeruginosa* and *S. aureus* attachment. The full set of topographical descriptors used for modelling is listed in **Table S1**, with a graphical illustration of selected topounits in **Fig S2**. The Random Forest machine learning regression algorithm was used to identify correlations between the topographic descriptors and bacterial attachment to the Topounits, generating high values for the coefficient of determination (R^2^) and low error values. Shapley Additive Explanations (SHAP) were used to identify the descriptors with the largest contributions to the Random Forest models.

The performance of the *P. aeruginosa* attachment model is shown in **Fig 2a,b**, and the results for *S. aureus* are shown in **Fig 2e,f**. The contributions of the five descriptors found to be the most important for each model are shown in **Fig 2c** for *P. aeruginosa* and in **Fig 2g** for *S. aureus*. The machine learning model predicted the observed attachment values for topographies in the test sets for both bacterial species models with high efficacy: R^2^ = 0.851±0.001 for *P. aeruginosa* average fluorescence and R^2^ = 0.810±0.001 for *S. aureus* average fluorescence (**Figs 2b and 2f**). The models were trained using the most significant surface parameters identified by the descriptor selection method SHAP. The most important descriptors for predicting bacterial attachment were the total area of the topographical features (feature coverage), the size of the spaces between objects (defined using the radii of inscribed circles illustrated in **Fig S1b**), and the radii of the maximum features. The topographical feature area was reflected in the model by a correlation with low bacterial attachment with both the pattern feature coverage and the maximum feature area within the unit cell in which the features are arranged. To extract design rules from the ML model, we plotted the individual top predictive descriptors in **Fig S4a** which indicate that the lowest *P. aeruginosa* attachment is associated with topographies with feature coverage > 0.5, maximum *inscribed circle radius* ≤ 3 µm, although unique solutions were not found for maximum feature radius alone. For *S. aureus*, the lowest attachment was associated with feature coverage > 0.4 and *maximum inscribed circle radius* ≤ 3 µm.

**Figure 2.**
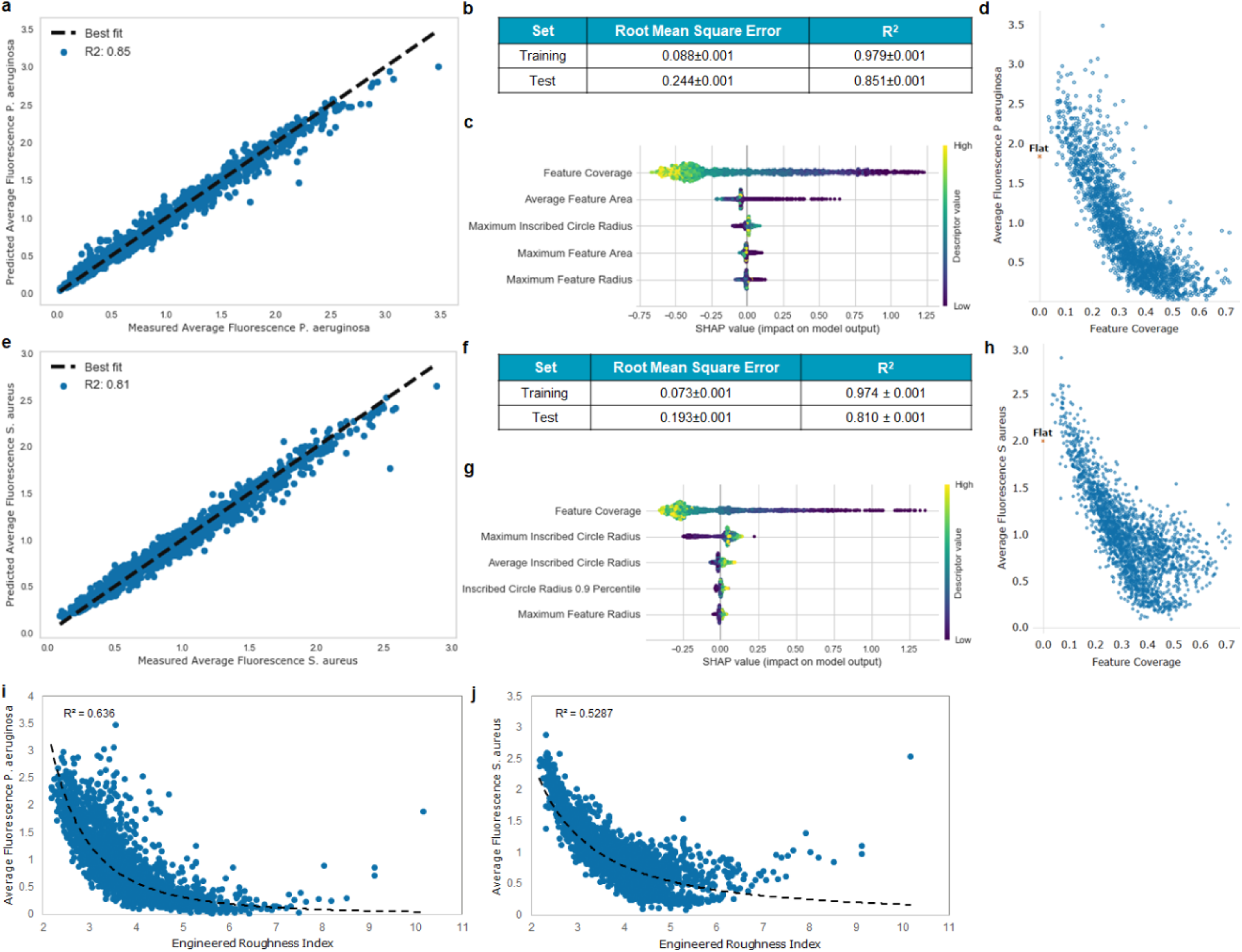
Machine learning modelling results for pathogen attachment using Random Forest: **(a)** Plot of the predicted versus measured average attachment values for one run of the *P. aeruginosa* test set, **(b)** Regression model performance metric results for *P. aeruginosa* training and test sets, **(c)** *P. aeruginosa* descriptors ranked by their importance from top to bottom and how their value (high = yellow or low = purple) impacts on the model (positive or negative impact), **(d)** Feature coverage descriptor distribution against attachment for *P. aeruginosa*, **(e)** Scatter plot of the measured against predicted average attachment values for one run of the *S. aureus* test set, **(f)** Regression model performance metric results for the *S. aureus* training and test sets, **(g)** *S. aureus* descriptor importance and their impact on model output. The topographical descriptors found to be most important for bacterial attachment are the inscribed circle which relates to the space between primitives, the total area covered by features (feature coverage) and the maximum feature radius, **(h)** Feature coverage descriptor distribution against attachment for *S. aureus*, **(i)**Comparison between P. *aeruginosa* attachment and the engineered roughness index (ERI) calculated for the topographies investigated (Schumacher et al^14^), **(j)** Comparison between S. *aureus* attachment and the roughness index calculated for the topographies.

The strong correlation of low attachment with the proportion of the surface covered with features and radius partnered with the negative correlation with the measurement of the trough dimensions (maximum inscribe circle radius), are powerful. These observations form the basis for design rules that can be used to develop biofilm resistant surfaces primarily describing the size of the elevated features (*feature coverage* and *maximum feature radius*) and the gaps between them (*maximum inscribed circle*).

Previously, the Engineered Roughness Index (ERI) calculated from micro topography dimensions, has been proposed by Schumacher et al^14^ to describe the decrease in microbial attachment for marine microorganisms (Ulva spore settlement) from a series of 4 pattern types and smooth silicone surfaces. The ERI encompasses three variables associated with the size, geometry, and spatial arrangement of the topographical features: Wenzel’s roughness factor (r) which is the ratio of the actual surface area to the projected planar surface area, multiplied by the depressed surface fraction (fD), and divided by the number of degrees of freedom for movement (d_f_). Notably, these elements are similar to the dominant descriptors in the model generated by ML from our topography library, i.e. parameters representing the area of features and the depressed areas between them. We calculated the ERI for all our topographies and plotted the relationship with bacterial attachment in **Figs 2i** and **2j** for *P. aeruginosa* and S. *aureus* respectively. It is clear that the trends observed confirm the hypothesis of Schumacher et al. that topographical patterns with higher roughness indexes reduce bacterial attachment and a non-linear relationship exists. However, the lower R^2^ values compared to our ML derived models (**Figs 2a and 2e**) indicate that the fit and therefore model is inferior likely due to the increased number of relevant topography description terms the ML model has identified.

The best performing micro topography from the Schumacher study was the bioinspired Sharklet pattern which they went on to test against *P. aeruginosa* and *S. aureus*. ^15^ Comparable wild-type *P. aeruginosa* data are not available on Sharklet, but for *S. aureus*, this surface in silicone was shown experimentally to provide a 67% reduction which is surpassed by our best performing topographies presented in **Fig 2h**.

### Micro topographical performance is independent of pathogen, growth environment and material

Based on the screening data obtained from quantification of *P. aeruginosa* and *S. aureus* attachment to polystyrene Topounits, those exhibiting a 5 to 20-fold reduction in bacterial attachment compared with a flat control surface (anti-attachment Topounits) were chosen for further study. To compare bacterial responses to these topographies with other patterned surfaces, pro-attachment Topounits with similar attachment to those of flat controls were selected (**Fig S2**). In **Fig 3**, representative images (**Fig 3a**) and quantitative data (**Fig 3b**) on selected Topounit scale-ups are presented for *P. aeruginosa, S. aureus, Proteus mirabilis* and *Acinetobacter baumannii* respectively grown under static conditions for 4 h. The data show that all four pathogens responded similarly to the polystyrene (PS) pro- (Topounits 336 and 697) and anti- (PS Topounits 685 and 881) attachment topographies.

**Figure 3.**
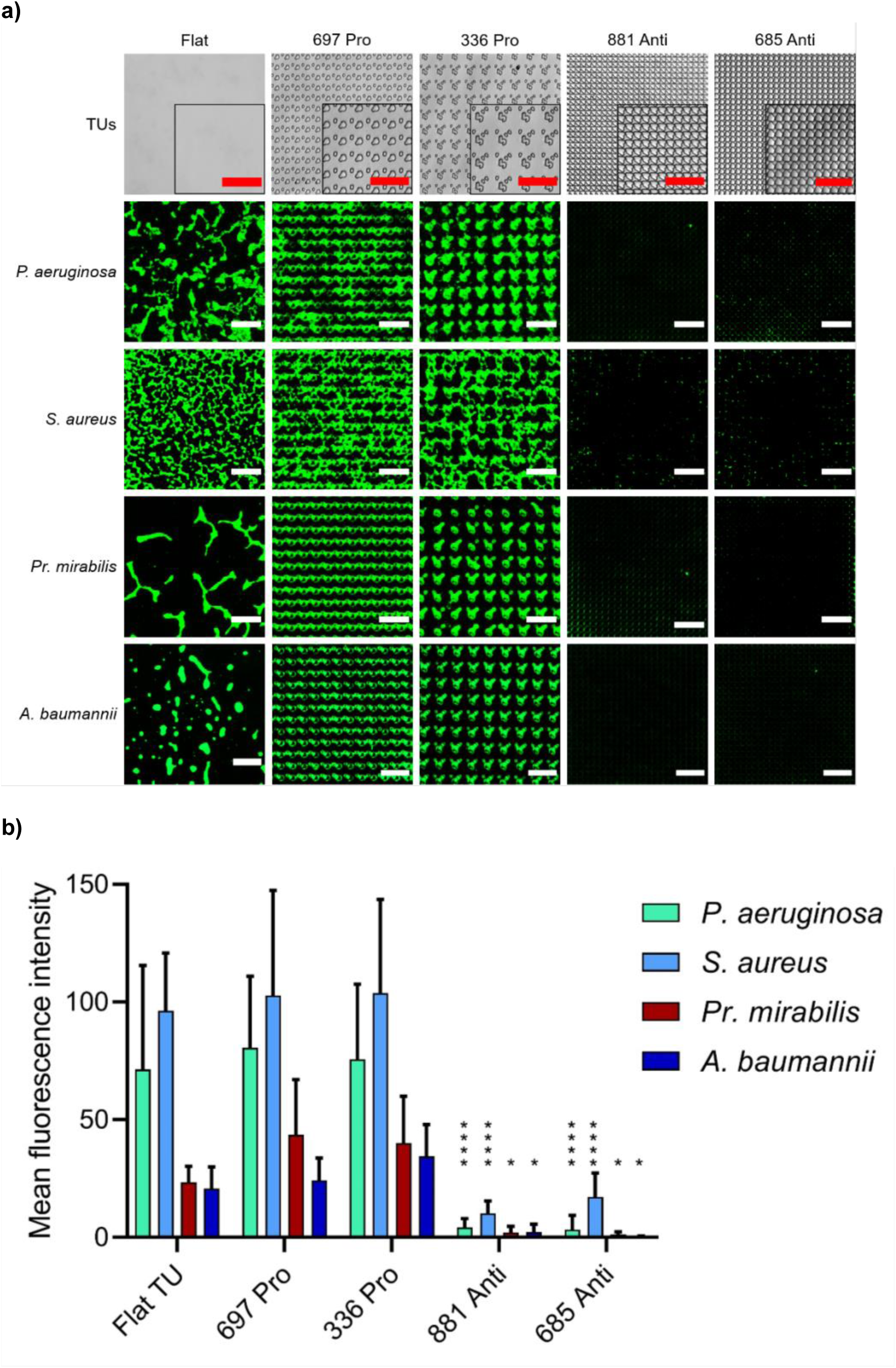
Attachment of bacterial pathogens to flat, pro- and anti-attachment polystyrene Topounits (**a**) Representative images of *P. aeruginosa, S. aureus, Pr. mirabilis* and *A. baumannii* attachment (bright field images of the Topounits shown in top row) after 4 h incubation under static conditions. (**b**) Quantification of mean fluorescence intensity for *P. aeruginosa, S. aureus, Pr. mirabilis* and *A. baumannii* cells stained with Syto9 and grown on the same Topounits. Scale bar: 50 µm. Data shown are mean ±SD, n ≥ 6. Statistical analysis was done using a two-way ANOVA with Dunnett’s multiple comparisons test (* *p*<0.05; ** *p*<0.01; *** *p*<0.001; **** *p*<0.0001).

We next investigated the performance of the Topounit scale-ups with respect to growth environment for motile *P. aeruginosa*. Inversion of the culture set up was used to assess the contribution of gravity to bacterial cell colonization. The key topographies maintained their *P. aeruginosa* pro- and anti-attachment properties independent of topographical orientation and also when flow was introduced (**Figs 4a and b**). The attachment of *P. aeruginosa* to two medically certified polymers, polyurethane (PU) and cyclic olefin copolymer (COC) embossed with the anti and pro-biofilm topographies was also assessed (**Fig 4a and b**). The selected micro topographies fabricated in both materials showed similar attachment levels to those in PS indicating that the surface topography strongly influences bacterial attachment independent of surface chemistry (**Figs 4a and b**).

**Figure 4.**
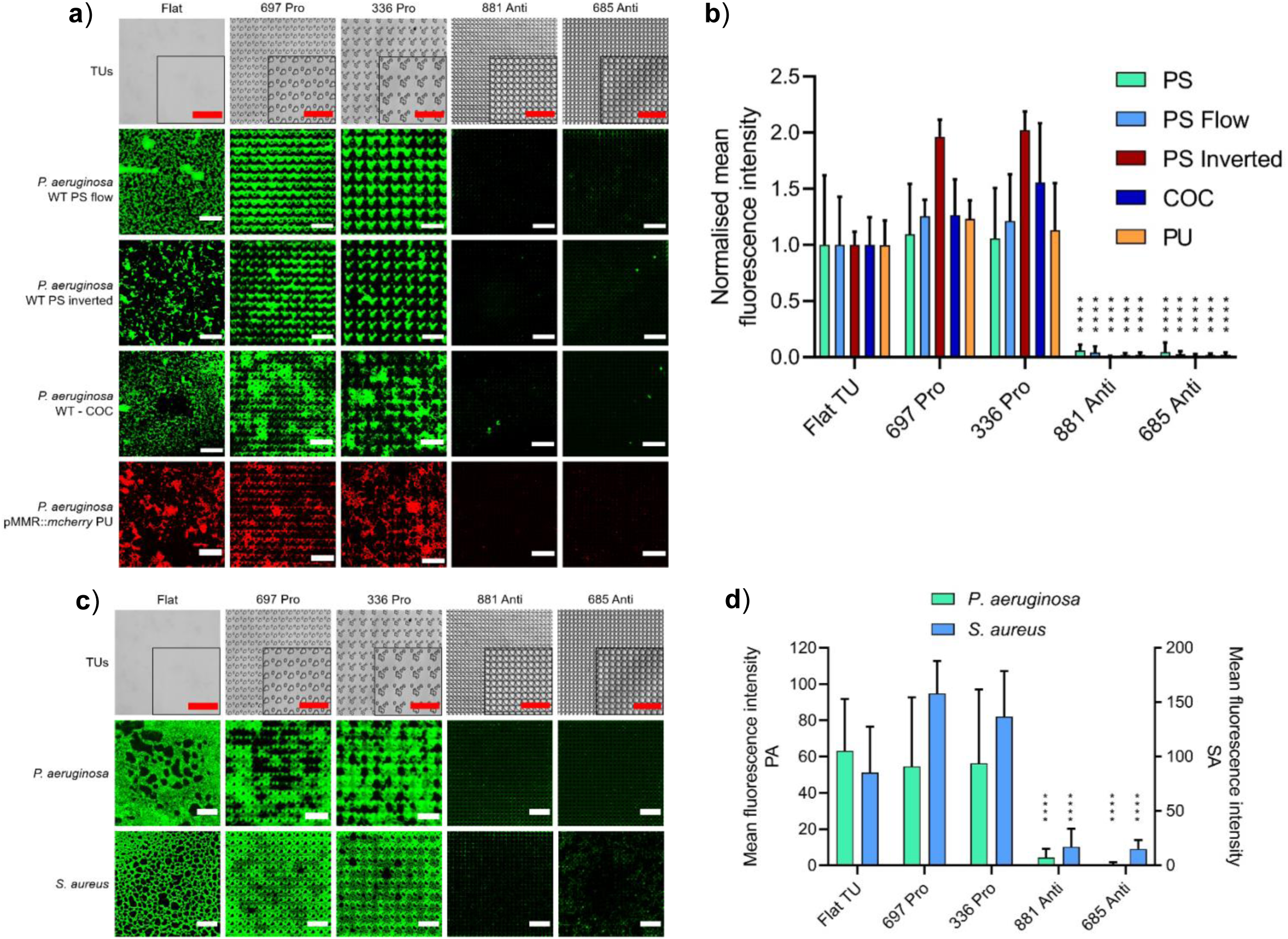
Bacterial attachment comparing different materials, inversion and flow. (**a**) Representative images of *P. aeruginosa* pME6032::*mcherry* (red) and *P. aeruginosa* wildtype stained with Syto9 fluorescent dye (green) grown for 4 h on flat, pro- or anti-attachment Topounits (bright field images shown in top row) moulded from polystyrene (PS), polyurethane (PU) or cyclic olefin copolymer (COC). *P. aeruginosa* wildtype attachment on upside down oriented Topounits (inverted) and under flow conditions is shown for polystyrene Topounits. (**b**) Quantification of mean normalised fluorescence intensity of *P. aeruginosa* incubated under conditions described above. (**c**) Bacterial biofilm assessment: Representative images of *P. aeruginosa* and *S. aureus* biofilms on flat, pro and anti-attachment PS topographies after 24 h incubation under static conditions. (**d**) Quantification of mean fluorescence intensity for *P. aeruginosa* and *S. aureus* biofilms shown in (**c**). Scale bar: 50 µm. Data shown are mean ±SD, n ≥ 5. Statistical analysis was done using a two-way ANOVA with Dunnett’s multiple comparisons test (* *p*<0.05; ** *p*<0.01; *** *p*<0.001; **** *p*<0.0001).

To determine whether *P. aeruginosa* and *S. aureus* attach and form biofilms on pro- and anti-attachment topographies after extended incubation, they were grown on PS Topounits for 24 h (**Fig 4c**). The anti-attachment PS Topounits 881 and 685 showed more than 15-fold lower *P. aeruginosa* biofilm formation compared to the flat surface (**Fig 4d**) (two-way ANOVA, *p*<0.0001). Mature biofilms incorporating interconnecting bacterial aggregates were clearly observed on the pro-attachment Topounits 697 and 336, as well as on the flat control, while the few cells present on the anti-attachment Topounits 881 and 685 were far more dispersed. Similar results were also obtained for *S. aureus* where a 5-fold (two-way ANOVA, *p*<0.0001) average reduction of surface growth on the two anti-attachment Topounits were observed after a 24 h incubation when compared with the flat control surface. Collectively, these data show that the 4h attachment data was a good predictor of subsequent biofilm resistance.

Remarkably, we observed that in these *in vitro* experiments with a motile Gram-negative organism (*P. aeruginosa*) and a non-motile Gram-positive organism (*S. aureus*) similar surface descriptors contributed to the attachment models for both species. The responses of *Pr. mirabilis* (motile) and *A. baumanni* (non-motile) to these selected pro- and anti-attachment Topounits were very similar to those of *P. aeruginosa* and *S. aureus*.

### Exploration of micro topographies by *P. aeruginosa*

Since bactericidal activity has been ascribed to various textured, mainly nano topographical features,^20-22^ we sought to determine whether a similar mechanism was responsible for the reduced colonisation of anti-attachment Topounits by *P. aeruginosa*. The viability of bacteria attached to the selected micropatterns was evaluated using live/dead differential cell staining. No significant differences were apparent in Live: Dead ratio for the selected topographies (**Fig S5**), indicating that these topographies did not exhibit bactericidal activity that would explain the differences in surface colonisation.

The outcome of bacterial exploration of a surface depends on the interplay between bacterial surface sensing and the local environment including surface physicochemical properties and architecture.^26^ The lack of colonization of the anti-attachment topographies may be a consequence of surface avoidance or detachment following initial attachment or trapping/confinement of individual cells in the troughs between the features. Since *P. aeruginosa* exhibits flagellar-mediated swimming and type IV pilus-mediated twitching motilities and utilizes both appendages in surface sensing^27,28,29^, we investigated the interactions of *P. aeruginosa* wild type and the isogenic flagella (Δ*fliC*) and pilus (Δ*pilA*) mutants with selected micro topographies.

Live tracking of wild type cells over the first 4 h (**Video 1**) revealed the presence of *P. aeruginosa* within the troughs between the features of an anti-attachment Topounit, whereas the pro-attachment surface was typified by more free movement of the cells about the surface between the much more widely spaced features, with preferential attachment of cells near the feature bases (**Video 2**). This is consistent with the fluorescent images observed in **Figs 3a** and **4a** where the pro-attachment features are decorated with the fluorescent cells preferentially compared to the flat areas between them. **Fig 5** shows that in common with the wild type, neither Δ*pilA* nor Δ*fliC* mutants colonized the anti-attachment Topounit although both mutants show reduced levels of attachment to both flat and the pro-attachment Topounit (**Fig 5b**). Thus, the loss of flagellar or type IV pilus does not render *P. aeruginosa* ‘blind’ to surface topography. While neither surface appendage was essential for colonization of the pro-attachment Topounit, the lack of type IV pili resulted in microcolony segregation (**Fig 5a**).

**Figure 5.**
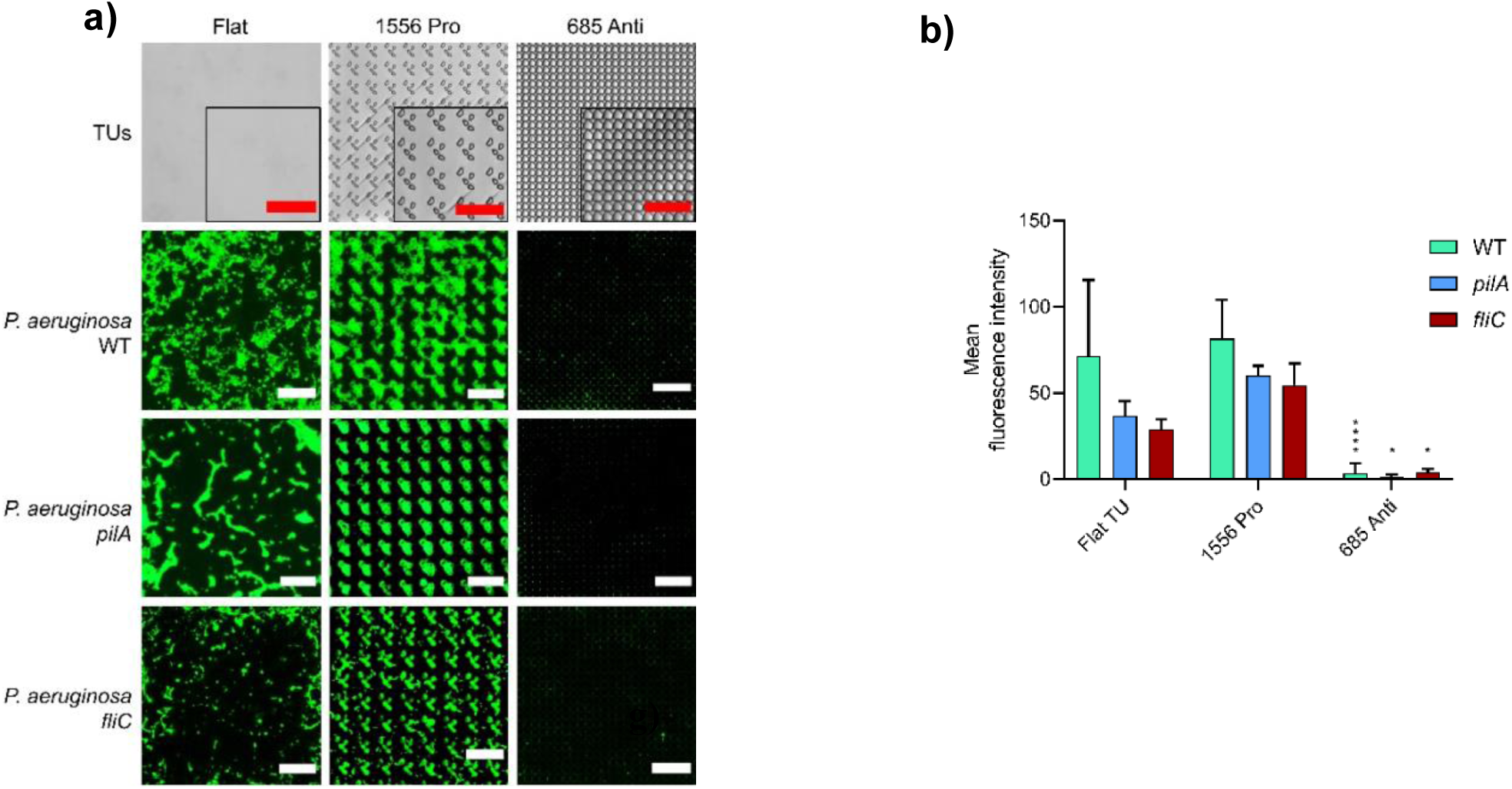
Early stage *P. aeruginosa* and *S. aureus* wildtype and *P. aeruginosa* motility mutant colonization of flat, pro- (1556) and anti-attachment (685) Topounits up to 4 h incubation in static conditions. (**a**) Representative fluorescent images showing *P. aeruginosa* wildtype, Δ*pilA* and Δ*fliC* mutants attached to each Topounits. (**b**) Mean fluorescence intensity of *P. aeruginosa* wildtype, Δ*pilA* and Δ*fliC* cells attached to each Topounits after 3.5 h.

The association of bacteria with troughs in substrates was previously observed for nanoscale patterned surfaces in work which identified cell localisation in nanoscale topographical features.^30^ Our observation that *P. aeruginosa* localise in the troughs between the raised features of the anti-attachment structures links to surface motility, with the rod-shaped cells having a greater propensity to be trapped at the bottom of high walled structures, something that has been reported for *P. aeruginosa* and modelled for engineered steps.^31^The differences in bacterial cell localisation for anti- and pro-attachment topographical patterns highlights the importance of early-stage cell confinement in the troughs between the structures, preventing later stage biofilm development. For *P. aeruginosa*, surface sensing and/or a failure to switch from reversible to irreversible attachment^27^ on the anti-attachment micro topographies are possible explanations for the differences the lack of attachment and subsequent biofilm formation observed.

### *In vivo* assessment of pro and anti-attachment topographies in a murine foreign body infection model

Prior to *in vivo* experiments it was important to determine whether surface conditioning by serum proteins influenced the interaction of bacterial cells with pro- and anti-attachment Topounits. Hence, we grew *P. aeruginosa* in tryptic soy broth (TSB) with or without human serum (10 % v/v) and compared protein adsorption to flat, pro- and anti-attachment Topounits after incubation at 37°C for 4h. Surface chemical analysis in **Fig S6** shows that serum protein treatment formed uniform layers (ToF SIMS) of similar thickness (XPS) were deposited on the pro- and anti-attachment Topounits. The relative attachment behaviours of *P. aeruginosa* on the flat, pro- and anti-attachment Topounits were not significantly affected by serum protein deposition (**Fig S7**).

To evaluate bacterial attachment resistance *in vivo*, scaled up (7 mm x 2 mm) Topounit features selected for their *in vitro* anti- or pro-attachment responses were embossed into polyurethane (PU) films and inserted subcutaneously in BALB/c mice. This polymer was used since it is sufficiently flexible not to irritate the mice and it is also a common biomaterial used for fabricating medical devices. After implantation, the mice were allowed to recover for 4 days and then inoculated with either *P. aeruginosa* (tagged with the fluorescent protein E2Crimson; **Table S2**) or PBS (uninfected control). After 8 days in the mice, the implants and surrounding tissues were recovered and sectioned. Retention of micro topographical feature integrity during the experiment was confirmed by scanning electron microscopy (**Fig S8**).

Mice were infected with *P. aeruginosa* PAO1-L expressing the fluorescent protein e2Crimson in order to visualise (**Fig 6a**) and quantify (**Fig 6b**) colonisation of the explant surface *in vivo*. The fluorescence obtained clearly demonstrate significantly lower colonisation of the anti-biofilm Topunit 881 compared with pro biofilm 697 and flatrespectively. Using a panel of markers (**Fig 6c**), migrating host cells labelled with nuclear stain DAPI (blue) or with the membrane dye FM1-43 (green) were observed to ingress into the Topounit regions in both the presence and absence of bacteria, penetrating the troughs between the features in the imaging plane shown. Since the Topochips were implanted 4 days prior to infection, it is likely that host cell colonization of the implants contributes to the reduction in bacterial attachment, in essence winning the ‘*race for the surface’*.^32^ Since bacterial attachment was prevented by anti-biofilm topographies *in vitro* in the absence of host cells, the intrinsic biofilm-resistant properties of specific Topounits such as 881 clearly play a role.

**Figure 6.**
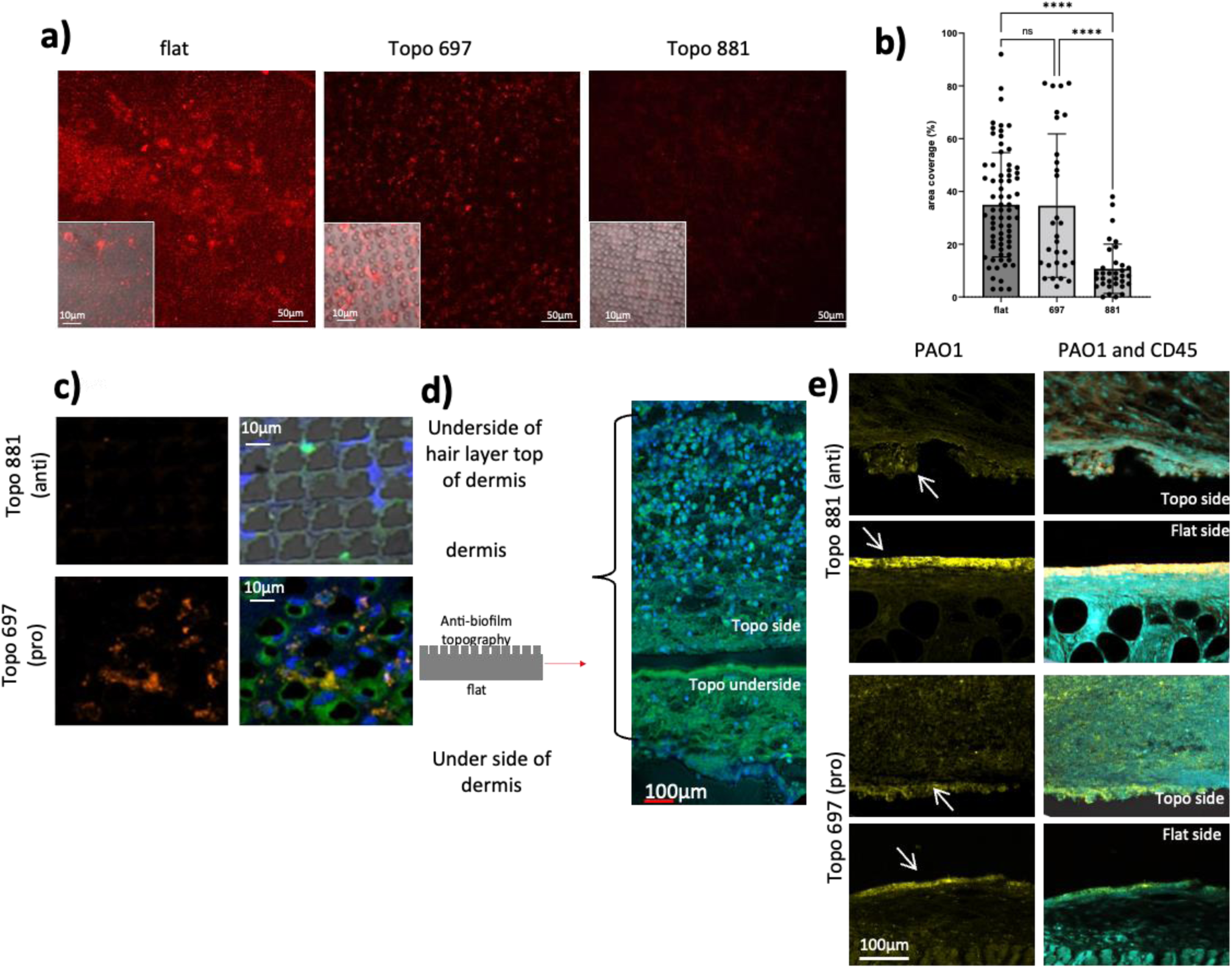
*In vivo* responses to anti- (881) and pro- attachment (697) Topounits implanted subcutaneously in mice. (**a**) *Red = P. aeruginosa* PAO1 expressing the fluorescent protein E2Crimson was used to quantify bacterial coverage on explanted Topounits 881 n=7, 697 n=8 and flat n=12. (**b**) Quantification of *Pseudomonas aeruginosa* from individual explants using constitutive E2Crimson fluorescence. (**c**) *Images of explanted Topounits:* orange = PAO1 bacteria detected using IHC antibody PA1-73116, secondary anti-rabbit Alexa 555. Overlay of blue = DAPI DNA stain green = FM1-43, cell membrane detection from both bacterial and host response and brightfield image. (**d**) **Tissue section** sterile experiment including **Topo side** increased host cell migration to surface; the flat **topo underneath**, showing asymmetrical host response to the implant. Blue = DAPI DNA stained nuclei, green = FM1-43 stained membranes. (**e**) **Tissue section** Infected with bacteria: overlay of asymmetrical tissue architecture that abutted the implant; **PAO1 & PAO1 and CD45**: bacterial localisation in tissues abutting the implant; yellow = *P. aeruginosa* PAO1 bacteria detected using IHC antibody PA1-73116, secondary anti-rabbit Alexa 555m (see white arrows), turquoise = CD45 cells from leukocyte lineage.

An asymmetrical host response to the Topounits, represented by staining of a representative polyurethane Topounit recovered from an un-infected (sham-inoculated) animal is shown in **Fig 6d**. The implant was removed from the mouse tissues during sectioning, leaving a void observed as a black area in each image. Host responses to the topographically patterned (top)_ and flat sides (bottom) were very different. Greater host cell migration (Dapi-stained nuclei, blue) into the Topounit side of the implant compared with the flat underside was apparent (**Fig 6d**). In **Fig 6e** (left-hand panel) an example of the differential distribution of bacteria on the patterned and flat sides of Topounits 881 and 697 is shown. *P. aeruginosa* (labelled yellow and highlighted with arrows) was primarily present in the tissue on the flat side of the implant. Host leukocyte cell infiltration is shown in **Fig. 6e** (righthand panel) as indicated by the CD45 positive cells (labelled turquoise). The importance of the design of topographically patterned polymers has been noted *in vitro* in terms of macrophage recruitment,^13^ but this, to our knowledge, is the first finding that that the host cell response *in vivo* is influenced by topography.

### Outlook

We have identified a range of topographical patterns and descriptors identified by machine learning that elicit *in vitro*, a reduction in bacterial attachment and subsequent biofilm formation for four different pathogens commonly associated with medical device associated infections. These findings could not have been anticipated from our current understanding of bacteria-surface interactions. Consequently, they illustrate the strength of unbiased screening of large topographical libraries to reveal previously unperceived cell–surface interactions, as well as providing insights towards the rational fabrication of new bioactive surfaces.^24,25^

*In vivo* experiments with *P. aeruginosa* indicate that these topographies recruit host cells and reduce infection. Texturing the surface of clinically approved biomaterials would benefit by retention of their bulk physical and mechanical properties while significantly reducing the need for expensive new materials discovery and commercial development. Combined with observations that material topography influences macrophage attachment and phenotype^13^, this work raises the exciting prospect of exploiting micro topographies to modulate host immune responses and prevent both biofilm-centred infections and foreign body reactions to implanted medical devices.

## ACKNOWLEDGEMENTS

This work was supported by the Engineering and Physical Sciences Research Council [grant nos. EP/N006615/1 and EP/K005138/1] the Wellcome Trust [grant nos. 103882 and 103884], the Biotechnology and Biological Sciences Research Council [BB/R012415/1], the Dutch Science Foundation (NWO) [grant VENI 15075], and the Dutch province of Limburg. We thank Dr Emily F Smith at the Nanoscale and Microscale Research Centre (NMRC - University of Nottingham) for acquiring XPS spectra and Dr Marta M Paino with XPS data interpretation. TopoChips were imaged in the School of Life Sciences Imaging Unit (SLIM - University of Nottingham) with help from Robert Markus and Seema Bagia. We thank Nick Beijer, and Nadia Roumans for TopoChip fabrication assistance, Chris Gell and Arsalan Latif for their help with data acquisition.

## AUTHORS’ CONTRIBUTIONS

MRA, JdB and PW conceived the project. MR and JL designed and conducted the in vitro and in vivo experiments respectively. JL, PW and AMG analysed in vivo data. GF, AuC, AV and DW conducted the modelling and the machine learning. SV and CB fabricated the TopoChip arrays, DS analysed the Topounits surface chemistry, JD constructed the mutants, AC and EI acquired and analysed the bacterial single cell tracking data. MR, MRA and PW wrote the manuscript with input from all other authors.

## COMPETING INTERESTS STATEMENT

We have no competing interests.

**VIDEOS** Files can be found here **https://bacterialweb.nottingham.ac.uk/**

Video 1: PAO1 on polystyrene anti -biofilm topography 685.

Video 2: PAO1 on polystyrene pro -biofilm topography 1556.

## METHODS

### Bacterial strains and culture conditions

Bacterial pathogens commonly associated with medical device-associated infections (Percival et al., 2015) chosen for this study included the Gram negatives, *P. aeruginosa* PAO1-L (Lausanne sub-line), *Pr. mirabilis* Hauser 1885 and *A. baumannii* ATCC17978 and the Gram positive, *S. aureus* SH1000 and are listed in **Table S2**. Bacteria were routinely grown at 37°C in lysogeny broth (LB) or LB agar supplemented with antibiotics as required. Tryptic soy broth (TSB) was used as the growth medium for bacterial TopoChip attachment assays. To mimic *in vivo* conditions, TSB supplemented with 10% v/v human serum (TSB HS10%) was used for some *P. aeruginosa* experiments. For polyurethane (PU) TopoChip attachment assays, *P. aeruginosa* PAO1-L carrying the constitutively expressed *mcherry* gene on the plasmid pMMR (Popat et al., 2012) was used to avoid the problem of PU autofluorescence.

### Construction of *P. aeruginosa* PAO1 flagella and type IV pili deletion mutants

*P. aeruginosa* PAO1 Δ*fliC* and Δ*pilA* mutants were constructed via 2-step allelic exchange obtained by double crossover as previously described (Hook et al., 2019). Two PCR products amplifying the upstream and the downstream nucleotide regions of each gene were generated using the primer pair 1FW/1RW and 2FW/2RW, respectively (**Table S1**). Both PCR products were fused by overlapping PCR to create a deletion in the corresponding gene. The resulting fragment was cloned into the suicide plasmid pME3087 (**Table S1**). The integration of the suicide plasmid into the *P. aeruginosa* chromosome was carried out by conjugation. Recombinants were selected on tetracycline (125 μg ml^-1^). The second cross over event was carried out using the carbenicillin enrichment method (300 μg ml^-1^) to select for the loss of tetracycline resistance by plating the resulting colonies on LB supplemented with or without tetracycline. The in-frame deletions were confirmed by PCR and DNA sequence analysis and their phenotypes confirmed by swimming and twitching motility assays (Morris et al., 2013; Hook et al., 2019).

### TopoChip fabrication

The TopoChip was designed by randomly selecting 2,176 features from a large *in silico* library of features containing single or multiple 10 μm high pillars within a virtual square of either 10 by 10, 20 by 20, or 28 by 28 μm^2^ size. (Unadkat et al, 2011) Micro-pillars were constructed from three different microscale primitive shapes: circles, triangles and rectangles (3 μm widths). Topographies were assembled as periodical repetitions of the features within 290 × 290 μm micro-wells (TopoUnits - Topounits) surrounded by 40 μm tall walls in a 66 × 66 array containing duplicate Topounits for each topography together with flat control surfaces. TopoChips were fabricated on a 2 × 2 cm^2^ chip divided into four quadrants as previously described.(Unadkat et al, 2011) Briefly, the inverse structure of the topographies was produced in silicon by standard photolithography and deep reactive etching. A mould fluorosilane release agent, trichloro (1H, 1H, 2H, 2H-perfluorooctyl) silane (Sigma Aldrich) was applied using vapour deposition. This silicon mould was then used to make a positive mould in poly(dimethylsiloxane) (PDMS) from which a second negative mould in OrmoStamp hybrid polymer (micro resist technology Gmbh) was fabricated. This served as the mould for hot embossing polystyrene (PS) (DSM), polyurethane (PU) (Elastollan, GoodFellow) and cyclic olefin copolymer (COC) GoodFellow films to produce TopoChips with 3 different chemistries. After fabrication the arrays were subjected to oxygen plasma etching. **Fig S1** shows examples of Topounits micro topographies. Each micro topography was assigned a feature index (FeatIdx) number (from 1 to 2176). [http://bacterialweb.nottingham.ac.uk/node_list2/21005/]

To ensure that the bacteria-material interactions observed were specifically dependent on surface topography, polystyrene TopoChips were analysed by time-of-flight secondary ion mass spectrometry (ToF-SIMS) together with X-ray photoelectron spectroscopy (XPS) for quantitative elemental analysis. A low level of mould release agent (F = 2 at%) was found to be transferred to the surface by the Topounit production process (**Fig S1a**) which was unavoidable due to the moulding requirements. It is clear from the chemical images that this is distributed unevenly according to the specific topography, which would be problematic since it would provide uneven chemistry from one Topounit to another, but the oxygen etching used initially to increase mammalian cell attachment was found to provide a uniform distribution within (**Fig S1b**) and between samples (**Fig S1c**). The Topochips were therefore deemed suitable for investigating response to the micro topographies since the while the chemistry was not pure oxygen etched polystyrene, it was the same on each unit.

### Surface chemical analysis

The surface chemistry of the TopoChips was assessed using time-of-flight secondary ion mass spectrometry (ToF-SIMS) and X-ray photoelectron spectroscopy (XPS). ToF-SIMS measurements were conducted on an ION-ToF IV instrument using a monoisotopic Bi_3_^+^ primary ion source operated at 25 kV and in ‘bunched mode’. TopoChip surface chemistry was quantified in terms of elemental composition using an Axis-Ultra XPS instrument (Kratos Analytical, UK) with a monochromated Al k*α* X-ray source (1486.6eV) operated at 10 mA emission current and 12 kV anode potential (120 W). Small spot aperture mode was used in magnetic lens mode (FoV2) to measure a sample area of approximately 110 µm^2^. CasaXPS (version 2.3.18dev1.0x) software was used for quantification and spectral modelling. The measured N 1s fraction for growth medium serum-conditioned surfaces was converted into protein layer thickness using Ray & Shard (2011) relationship between [N] and protein depth.

### TopoChip screen for bacterial attachment and viability

Prior to incubation with bacteria, TopoChips were washed by dipping in distilled water and sterilized in 70% v/v ethanol. The air-dried chips were placed in petri dishes (60mm x 13mm) and incubated statically or with shaking (60 rpm) at 37°C in 10 ml of growth medium inoculated with diluted (optical density: OD_600 nm_ = 0.01) bacteria from overnight cultures. Static incubation was selected for screening experiments as this produced consistent bacterial attachment while reducing the formation of biofilm streamers on TopoChip corners which can induce cross contamination of Topounits under flow (Rusconi et al., 2010). At specific time points, TopoChips were removed and washed in phosphate buffered saline (PBS) pH 7.4 to remove loosely attached cells. After rinsing with distilled water, attached cells were stained with Syto9 (50 µM; Molecular Probes, Life Technologies) for 30 min at room temperature. After staining, TopoChips were rinsed with distilled water, air-dried and mounted on a glass slide using Prolong antifade reagent (Life Technologies). The viability of attached cells was evaluated by fluorescent staining with the LIVE/DEAD® BacLight™ bacterial viability kit (Molecular Probes, Life Technologies).

### TopoChip imaging and data acquisition

TopoChips were imaged using a Zeiss Axio Observer Z1 microscope (Carl Zeiss, Germany) equipped with a Hamamatsu Flash 4.0 CMOS camera and a motorized stage for automated acquisition. A total of 4,356 images (one per Topounits) were acquired for each chip using a 488 nm laser as light source. Since the bacterial cells may attach at different heights on the micro-patterns, images were initially acquired as 50 µm range Z-stacks (2 µm steps - 25 slides) from the Topounits using a 40× objective (Zeiss, LD Plan-Neofluar 40×/0.6 Korr Ph 2 M27). Although this method facilitates capture of the total fluorescence emitted by bacteria attached to the topographies, it significantly increased scanning times and file sizes (∼460 GB per chip). Hence, a lower ×10 magnification lens (Zeiss, EC Plan-Neofluar 10×/0.30 Ph 1) was routinely used as it provided sufficient depth resolution for capturing the total fluorescence per Topounits. This enabled the use of the auto-focus function, which considerably reduced scanning times and file sizes per TopoChip. Cropping the images into a smaller field of view (247 µm x 247 µm) and omitting the walls of the micro-wells reduced the artefacts arising from bacterial attachment to the walls and further improved the auto-focus function.

To identify out-of-focus images from each TopoChip dataset, individual topographical images were combined into composites using open source Fiji-ImageJ 1.52p software (National Institutes of Health, US). To improve TopoChip dataset quality, image pre-processing also included a) staining artefact removal by excluding pixels with fluorescence intensities higher than 63,000 from data acquisition and b) assay-specific background fluorescence was removed using flat control Topounits as a reference.

To classify topographies influencing bacterial attachment, the mean fluorescence intensity on each Topounits was measured using Fiji-ImageJ. Hit micro topographies with anti-attachment properties were selected for further studies based on the screening data obtained from quantifying *P. aeruginosa* (n = 22) and *S. aureus* (n = 6) attachment to polystyrene TopoChips (**Fig S1**).

Additionally, and to discriminate between bacterial responses to hit topographies with other patterned surfaces, Topounits with similar or a small increase in attachment with respect to flat controls (“pro-attachment Topounits ”) were also selected for further investigation. To ascertain that hit surfaces influenced the observed bacterial responses, two-way ANOVA with Dunnett’s multiple comparisons tests were applied to determine whether bacterial attachment on Topounits differed significantly from that of a flat control (*p*<0.05) when compared with the variations within the replicates using GraphPad Prism 8.0 (GraphPad Software, Inc., San Diego, CA).

### Generation of predictive surface topography attachment models

To define the surface design parameters that influence bacterial attachment, the data acquired from screening *P. aeruginosa* and *S. aureus* on polystyrene TopoChips were interrogated as follows. The fluorescence values for bacterial attachment to each of the replicate Topounits were normalized to the average fluorescence intensity of the chip quadrant to account for differences in staining intensities between experiments. The fluorescence intensity was established to correlate with the number of attached fluorescent bacteria, as shown previously (Hook et al., 2012). It was therefore used as the dependent variable in the models. Topounits with low signal to noise ratio (< 2) were carefully excluded from the datasets of *P. aeruginosa* (77 units removed) and *S. aureus* attachment (10 units removed). The Random Forest machine learning method was applied to generate relationships between the topographies and bacterial attachment using the topographical descriptors listed in **Table S2**. The Random Forest module was used with default parameters in Python 3.7. Bootstrapping without replacement was used to define training and test sets for 50 model runs. Seventy percent of each bacterial attachment dataset was used to train the model, and 30% were kept aside to determine the predictive power of the model. The SHAP (SHapley Additive exPlanations) package in Python 3.7 was used to eliminate less informative descriptors and to determine descriptor importance.

### Murine foreign body infection model

To investigate the progress of bacterial infection and the host immune response to pro- (FeatIdx 697) and anti- (FeatIdx 881) attachment Topounits, a murine foreign body infection model was used (Hook *et al*, 2012). Scaled up Topounit micro-patterns were imprinted on one side of PU sheet of dimensions 3 mm x 7 mm. The Topunits or flat PU with the same dimensions were implanted subcutaneously using a 9-gauge trocar needle (1 per mouse, 3 repeats for each micropattern topography and flat), into 19–22 g female BALB/c mice (Charles River) with the patterned side facing upwards to the skin surface. One hour before implantation, 2.5 mg/kg of Rymadil analgesic (Pfizer) was administered by subcutaneous injection. Animals were anaesthetized using isoflurane, the hair on one flank removed by shaving and the area sterilized with Hydrex Clear (Ecolab). After foreign body insertion, mice were allowed to recover for 4 days prior to subcutaneous injection of either 1 × 10^5^ colony forming units (CFUs) of *P. aeruginosa* or vehicle (phosphate buffered saline; uninfected control).

Mice were housed in groups of 4 in ventilated cages under a 12 h light cycle, with food and water *ad libitum*, and with weight and clinical condition of the animals recorded daily. Four days post infection, the mice were humanely killed and the micropatterned PU Topounits samples and the surrounding tissues removed. PU Topounits samples were fixed in 10% v/v formal saline and labelled with antibodies targeting CD45 (pan-leukocyte marker; VWR violetfluor 450), and the membrane-selective dye FM1-43 (Thermofisher Scientific) for total Topounits-associated biomass. The distribution of bacterial cells on the micropatterned surfaces was visualized using two separate methods. Initially with polyclonal antibodies raised against *P. aeruginosa* (Thermofisher Scientific). Secondary antibodies used were goat anti-rabbit (Alexa 555; Thermofisher Scientific). Later *Pseudomonas aeruginosa* PAO1 Lausanne E2Crimson fluorescence was used to quantify bacterial coverage on the surface of explanted devices. Images acquired on a Zeiss 700 confocal microscope were quantified using Image J (image processing pipeline provided in supporting data). All animal work was approved following local ethical review at University of Nottingham and performed under U.K. Home Office Project License 30/3082.

## Notes

https://bacterialweb.nottingham.ac.uk/node_list2/25365/

### Competing Interest Statement

The authors have declared no competing interest.

### Summary of Updates

-Author list updated and new in vivo data included -link to online materials added

