## Supplementary Information for "Micro topographical instruction of bacterial attachment, biofilm formation and *in vivo* host response"

#### 1. Machine Learning Data Analysis

Software used to classify mammalian cell shape (Cell Profiler) was used to analyse the images used to create the Topounits, providing 66 uncorrelated topographical shape descriptors that were used to train *P. aeruginosa* and *S. aureus* attachment models. The full set of topographical descriptors is listed in **Table S1**.

For *P. aeruginosa,* bacterial attachment to 1,852 TUs were investigated and for *S aureus* 2,084 were considered (**Fig 1**). TUs were excluded from the analysis if their signal to noise ratio was lower than 2. The Random Forest machine learning method and Multiple Linear Regression with Expectation Maximisation (MLREM)(Burden and Winkler, 2009) were both used to generate non-linear and linear relationships between the topographies and bacterial attachment, producing good models for the datasets. Those methods were coupled with Shappley Additive Explanation (SHAP)(Lundberg and Lee, 2017) method for descriptor selection. Bootstrapping without replacement using 50 runs was used to assess the model robustness over multiple data samples. The models were built based on the top three most informative descriptors, common for both datasets, as identified by SHAP. All methods were implemented in Python 3.7. Random Forest from scikt-learn version 0.22.1 using default parameters was employed to generate the ML models. Seventy percent of each dataset was used to train the models, and 30% were kept aside in a test set used to determine the predictive power of the models.

Although Random Forest has produced a better non-linear fit to the data (with average $R^2$ = 0.85±0.001 and average RMSE 0.24±0.001 log fluorescence for *P. aeruginosa;* and average $R^2$ = 0.81±0.001 and RMSE 0.19±0.001 log fluorescence for *S. aureus* in the test set), MLREM regression coefficients assisted informing the individual contribution of each descriptor to attachment (**Figs 2B and 2F**). MLREM results showed that there is also a strong linear correlation between the selected descriptors and bacterial attachment, with $R^2$ = 0.69±0.016 and RMSE 0.35±0.012 log fluorescence for *P. aeruginosa;* and $R^2$ = 0.69±0.029 and RMSE 0.26±0.13 log fluorescence for *S. aureus* in the test set.

**Table S1**: TopoUnit topographical surface descriptors derived from image analysis of the photo lithography design files using CellProfiler and Image J.

| | |
|---|---|
| **Total Circle Area Scaled** | **The total area of circle primitives scaled by feature area [0,1].** |
| **Number of Colour Changes Diagonally** | Number of colour changes of the feature over the diagonal. |
| **Circle Area** | The sum of the area of circle primitives in pixels. |
| **Circle Diameter** | The sum of diameter of circle primitives in pixels. |
| **Number of Rectangles Scaled** | The number of rectangle primitives scaled by feature area [0,1]. |
| **Number of Triangles Scaled** | The number of triangle primitives scaled by feature area [0,1]. |
| **Feature Unit Cell Size** | Topo unit cells have been fabricated using 3 unit cell sizes within which each feature is placed: 10 x 10 µm, 20 x 20 µm and 28 x 28 µm. This descriptor indicates the size of the unit cell for the topo unit. |
| **Inscribed Circle Number** | Number of inscribed circles that can be inserted between features, as determined by ImageJ (See Figure S1b). |
| **Inscribed Circle Radius 0.1 Percentile** | 0.1 percentile values calculated for inscribed circles radii within a topo chip. |
| **Inscribed Circle Radius 0.25 Percentile** | 0.25 percentile values calculated for inscribed circles radii within a topo chip. |
| **Inscribed Circle Radius 0.75 Percentile** | 0.75 percentile values calculated for inscribed circles radii within a topo chip. |
| **Inscribed Circle Radius 0.9 Percentile** | 0.9 percentile values calculated for inscribed circles radii within a topo chip. |
| **MAD Inscribed Circle Radius** | Median Absolute Deviation of the inscribed circle radii in the topo chip. |
| **Maximum Inscribed Circle Radius** | Maximum value calculated for the inscribed circle radii in the topo chip. |
| **Average Inscribed Circle Radius** | Average value calculated for the inscribed circle radii in the topo chip. |
| **Median Inscribed Circle Radius** | Median value calculated for the inscribed circle radii in the topo chip. |
| **Minimum Inscribed Circle Radius** | Minimum value calculated for the inscribed circle radii in the topo chip. |
| **Mode Inscribed Circle Radius** | Model value calculated for the inscribed circle radii in the topo chip. |
| **Std Dev Inscribed Circle Radius** | Standard deviation value calculated for the inscribed circle radii in the topo chip. |
| **Rectangle Area** | The area of rectangular primitives. |
| **Rectangle Length** | The length of the rectangular primitives. |
| **Number of Rectangles** | Number of Rectangles in the topo unit cell |
| **Number of Triangles** | Number of Triangles in the topo unit cell |
| **Maximum Feature Area** | Area of the biggest feature in the topo unit cell. |
| **Average Feature Area** | Average feature area in the topo unit cell. |
| **Minimum Feature Area** | Minimum feature area in the topo unit cell. |

| | |
|---|---|
| **Maximum Feature Compactness** | The maximum value for compactness for the features in the topo unit cell. Compactness is calculated as the variance of the radial distance of the object's pixels from the centroid divided by the area. |
| **Average Feature Compactness** | The average compactness calculated for the features in the topo unit cell. |
| **Mode Feature Compactness** | The mode value for compactness for the features in the topo unit cell. |
| **Feature Compactness Percentile 0.1** | 0.1 percentile values for compactness calculated for the features in the topo unit cell. |
| **Skewness Feature Compactness** | Skewness of the variance of the radial distance of the object's pixels from the centroid divided by the area. |
| **Variance Feature Compactness** | Variance of the feature compactness values. |
| **MAD Feature Eccentricity** | Median absolute deviation of the feature's eccentricity. The eccentricity of the ellipse that has the same second-moments as the region. The eccentricity is the ratio of the distance between the foci of the ellipse and its major axis length. The value is between 0 and 1. (0 and 1 are degenerate cases; an ellipse whose eccentricity is 0 is actually a circle, while an ellipse whose eccentricity is 1 is a line segment.). |
| **Maximum Feature Eccentricity** | Maximum value for feature eccentricity in the topo unit cell. |
| **Average Feature Eccentricity** | Average value for feature eccentricity in the topo unit cell. |
| **Median Feature Eccentricity** | Median value for feature eccentricity in the topo unit cell |
| **Feature Eccentricity Percentile 0.1** | 0.1 percentile eccentricity values calculated for features in the topo unit cell. |
| **Skewness Feature Eccentricity** | Skewness of the eccentricity calculated for features in the topo unit cell. |
| **Variance Feature Eccentricity** | Variance of the feature eccentricity values. |
| **Maximum Feature Extent** | Maximum extent values for the features in a topo unit cell. Extend is the proportion of the pixels in the bounding box that are also in the region. Computed as the Area divided by the area of the bounding box. |
| **Average Feature Extent** | Average extent values for the features in a topo unit cell. |
| **Mode Feature Extent** | Mode extent values for the features in a topo unit cell. |
| **Feature Extent Percentile 0.1** | 0.1 percentile extent values calculated for the features in a topo unit cell. |
| **Variance Feature Extent** | Variance of the extent values for the features in a topo unit cell. |
| **Maximum Feature Form Factor** | Maximum form factor value calculated for the features in the topo unit cell. Form factors is calculated as $4*\pi*Area/Perimeter^2$. Equals 1 for a perfectly circular object. |
| **Average Feature Form Factor** | Average form factor value calculated for the features in the topo unit cell. |
| **Mode Feature Form Factor** | Mode form factor value calculated for the features in the topo unit cell. |
| **Variance Feature Form Factor** | Variance form factor value calculated for the features in the topo unit cell. |
| **MAD Feature Major Axis Length** | Median absolute deviation of the feature's major axis length. The length (in pixels) of the major axis of the ellipse |

| | |
|---|---|
| | that has the same normalised second central moments as the region. |
| **Maximum Feature Radius** | The maximum radius of the biggest feature in the topo unit cell. The radius is calculated as the distance of any pixel in the object to the closest pixel outside of the object. For high aspect ratio objects, this is 1/2 of the maximum width of the object. |
| **Average Feature Radius** | Average value calculated for the maximum radius of the features in the topo unit cell. |
| **Variance Feature Radius** | Variance value calculated for the maximum radius of the features in the topo unit cell. |
| **Variance Feature Minimum Feret Diameter** | The Feret diameter is the distance between two parallel lines tangent on either side of the object (imagine taking a calliper and measuring the object at various angles). The minimum Feret diameter is the smallest possible diameter, rotating the callipers along all possible angles. |
| **Maximum Feature Orientation** | Maximum orientation calculated for the features in the topo unit cell. Orientation is defined as the angle (in degrees ranging from -90 to 90 degrees) between the x-axis and the major axis of the ellipse that has the same second-moments as the region. |
| **Average Feature Orientation** | Average orientation calculated for the features in the topo unit cell. |
| **Median Feature Orientation** | Median orientation calculated for the features in the topo unit cell. |
| **Mode Feature Orientation** | Mode orientation calculated for the features in the topo unit cell. |
| **Feature Orientation Percentile 0.1** | 0.1 percentile orientation values calculated for the features in the topo unit cell. |
| **Variance Feature Orientation** | Variance orientation calculated for the features in the topo unit cell. |
| **Number of Features** | Number of features inside a topo unit cell. |
| **Std Dev of Rotation** | The standard deviation (in degrees), is used to determine the rotation of the primitives when they are placed in the feature. |
| **Total Area Triangle** | The total area of circle primitives scaled by feature area [0,1]. |
| **Feature Coverage** | Percentage of the total area occupied by the features in the topo unit. |
| **Total Perimeter** | Total perimeter of the features in the topo unit cell. |
| **Triangle Area** | The area of triangle primitives in the topo unit cell. |
| **Triangle Size** | Length of the shortest side of a triangle primitive in the topo unit cell. |
| **Percentage of Pixels Covered by Topographies** | Total number of pixels representing the topographies in the design in relation to the whole design image. This descriptor contains the same information as Total Area. |

**Table S2**: Bacterial strains, plasmids and primers used in this study

| Strain, plasmid or primer | Genotype and/or Relevant characteristic | Source or reference |
|---|---|---|
| **Strain** | | |
| *P. aeruginosa* | | |
| PAO1-L | Wild type PAO1 strain, Lausanne subline | B. Holloway via D. Haas |
| PAO1-W | Wild-type PAO1 strain, Washington subline | Washington collection |
| PAJD431 | In frame deletion of *pilA* in PAO1-W | This study |
| PAJD477 | In frame deletion of *fliC* in PAO1-W | This study |
| *S. aureus* | | |
| SH1000 | Wild-type | Horsburgh et al., 2002 |
| *Pr. mirabilis* | | |
| Hauser 1885 | Wild-type | Hauser, 1885 |
| *A. baumannii* | | |
| ATCC17978 | Wild-type | Baumann et al., 1968 |
| *E. coli* | | |
| DH5α | *recA1 endA1 hsdR17 supE44 thi-1 gyrA96 relA1 Δ(lacZYA-argF)U169[ϕ80 dlacZΔM15],* Nal$^R$ | Liss, 1987 |
| S17.1λpir | *thi pro hsdR hsdM*$^+$ *recA RP4-2-Tc::Mu-Km::Tn7 λpir*, Gm$^R$ | Simon *et al.*, 1983 |
| **Plasmids** | | |
| pME3087 | Suicide vector for homologous recombination, ColE1 replicon, Mob; Tc$^R$ | Voisard *et al.*, 1994 |
| pJD112 | pME3087 derivative for the generation of *pilA* in frame deletion mutant; Tc$^R$. | This study |
| pJD113 | pME3087 derivative for the generation of *fliC* in frame deletion mutant; Tc$^R$. | This study |
| P*cdrA*::*gfp*$^S$ | pUCP22Not-P*cdrA*-RBS-CDS-RNaseIII-*gfp*(Mut3)-T0-T1, Ap$^R$ Gm$^R$; c-di-GMP reporter | Rybtke *et a*l 2012 |
| **pSW002-PcE2-Crimson** | TcR Broad host-range expression vector with constitutive Pc promoter; for tagging bacteria with E2-Crimson | pSW002-PcE2-Crimson |
| PilAΔ FW1 | 5′-ATATCTAGAATGCCGAACTGCTCG-3′ | This study |
| PilAΔ RV1 | 5′-TTAGTTATCACAACCTTGAGCTTTCATGAATCTCTC-3′ | This study |
| PilAΔ FW2 | 5′-TTCATGAAAGCTCAAGGTTGTGATAACTAAGGTGAT-3′ | This study |
| PilAΔ RV2 | 5′-TATCTGCAGAAGTGGAAGTGGAGA-3′ | This study |
| FliCΔ FW1 | 5′-ATATCTAGAATGCTCGAAGGCGCGCATCT-3′ | This study |
| FliCΔ RV1 | 5′-TTAGCGCAGCAGGCTTGTAAGGGCCATGGTGATTTC-3′ | This study |
| FliCΔ FW2 | 5′-ACCATGGCCCTTACAAGCCTGCTGCGCTAAGCCCGG-3′ | This study |
| FliCΔ RV2 | 5′-TATAAGCTTAAGTCGTTCAACCCGCGCGT-3′ | This study |

### SUPPLEMENTARY FIGURES

**a)**

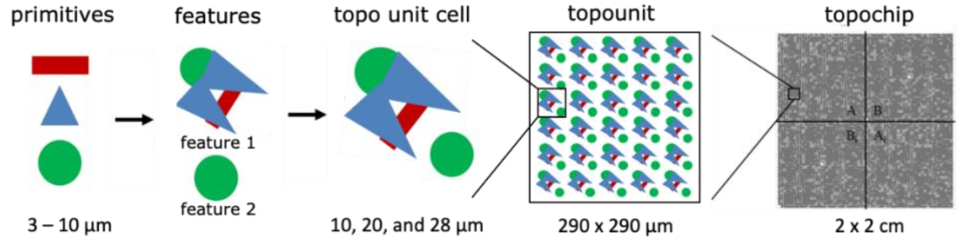


**b)**

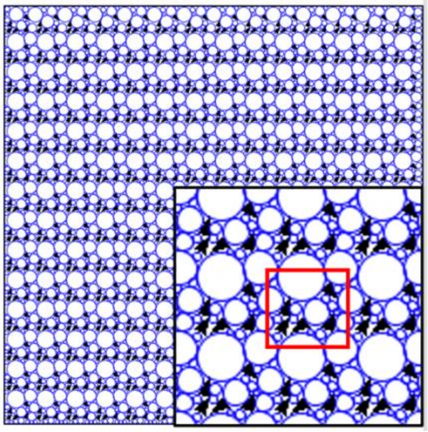

**Figure S1 a)** In the TopoChip, each micro topographical element contains primitives (circles, triangles and rectangles) which form features that are repeated to cover the surface of a TopoUnit within a unit cell with a size of either 10 x 10, 20 x 20, or 28 x 28µm.
**b)** Image analysis of example topounits, illustrating the inscribed circles (blue) used to describe the areas between the topo features for 6 different patterns (black). The unit cell is represented in red.

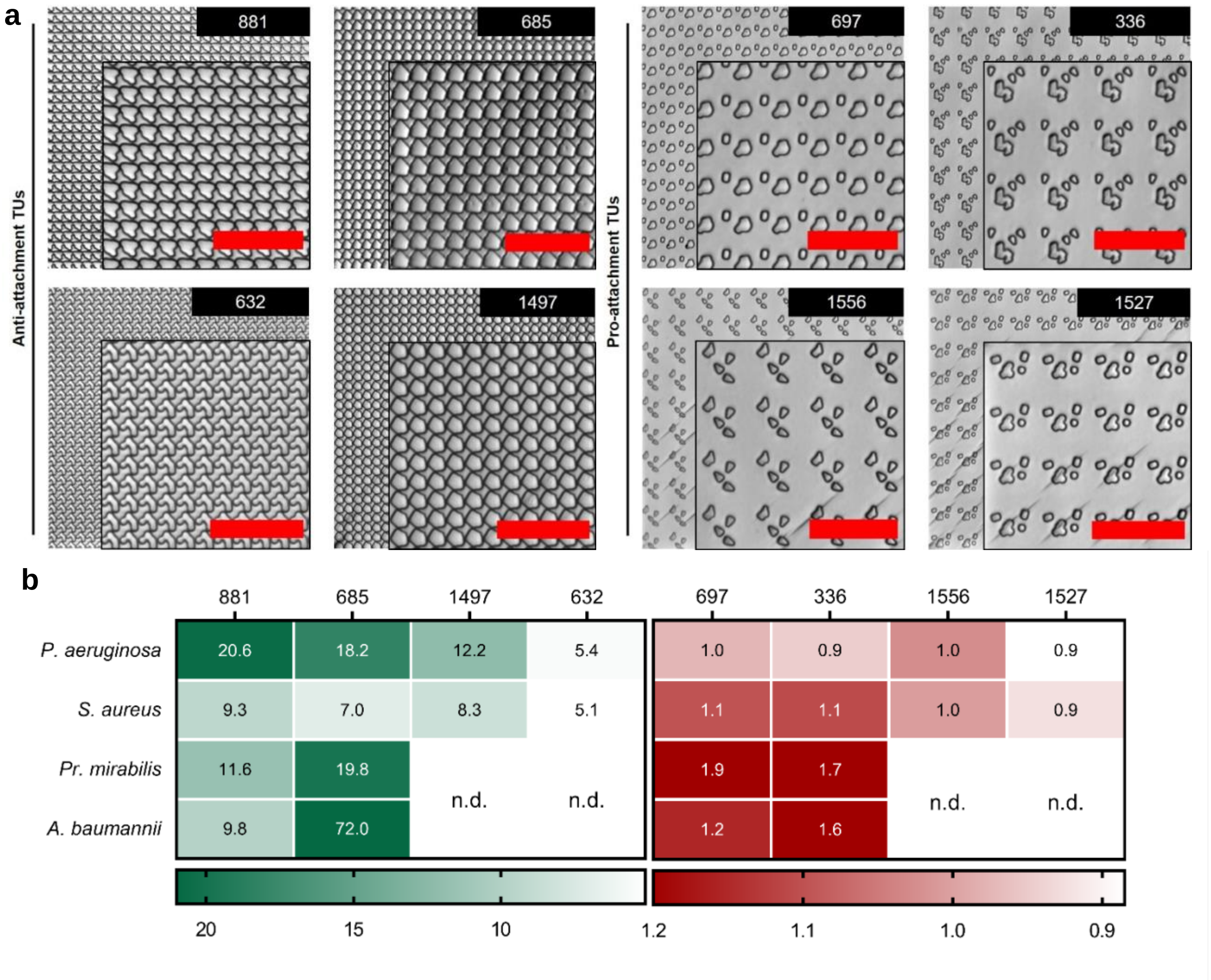


**Figure S2.** Selected anti- and pro-attachment micro topographies (**a**) based on the screening data obtained from quantifying *P. aeruginosa* and *S. aureus* attachment to PS TopoChips. (**b**) Intensity maps of the fold reduction (red) or increase (green) in the measured fluorescence of the flat control for *P. aeruginosa*, *S. aureus*, *Pr. mirabilis* and *A. baumannii* attachment to the same PS TUs after 4 h incubation in static conditions. Scale bar: 50 µm.

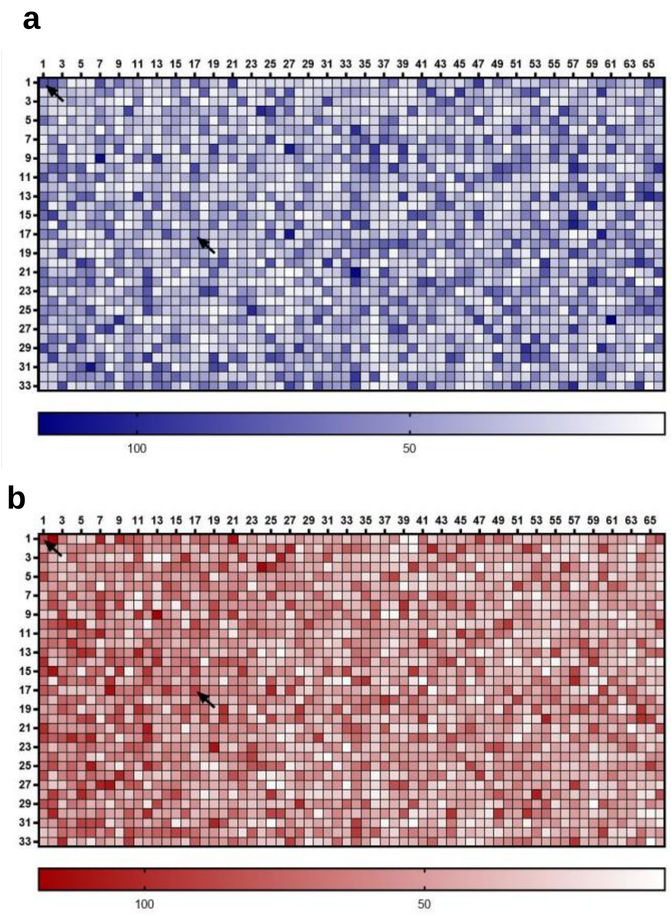


**Figure S3.** Intensity maps of measured fluorescence of *P. aeruginosa* (**a**) and *S. aureus* (**b**) attached to TUs from PS TopoChips after 4-h incubation in TSB + HS10% or TSB respectively. Shading within each outlined square indicates the mean fluorescence intensity value for the TU (Key to bottom). Black arrows pointing towards the TU coordinates 1,1 and 17,17 indicate flat surface controls.

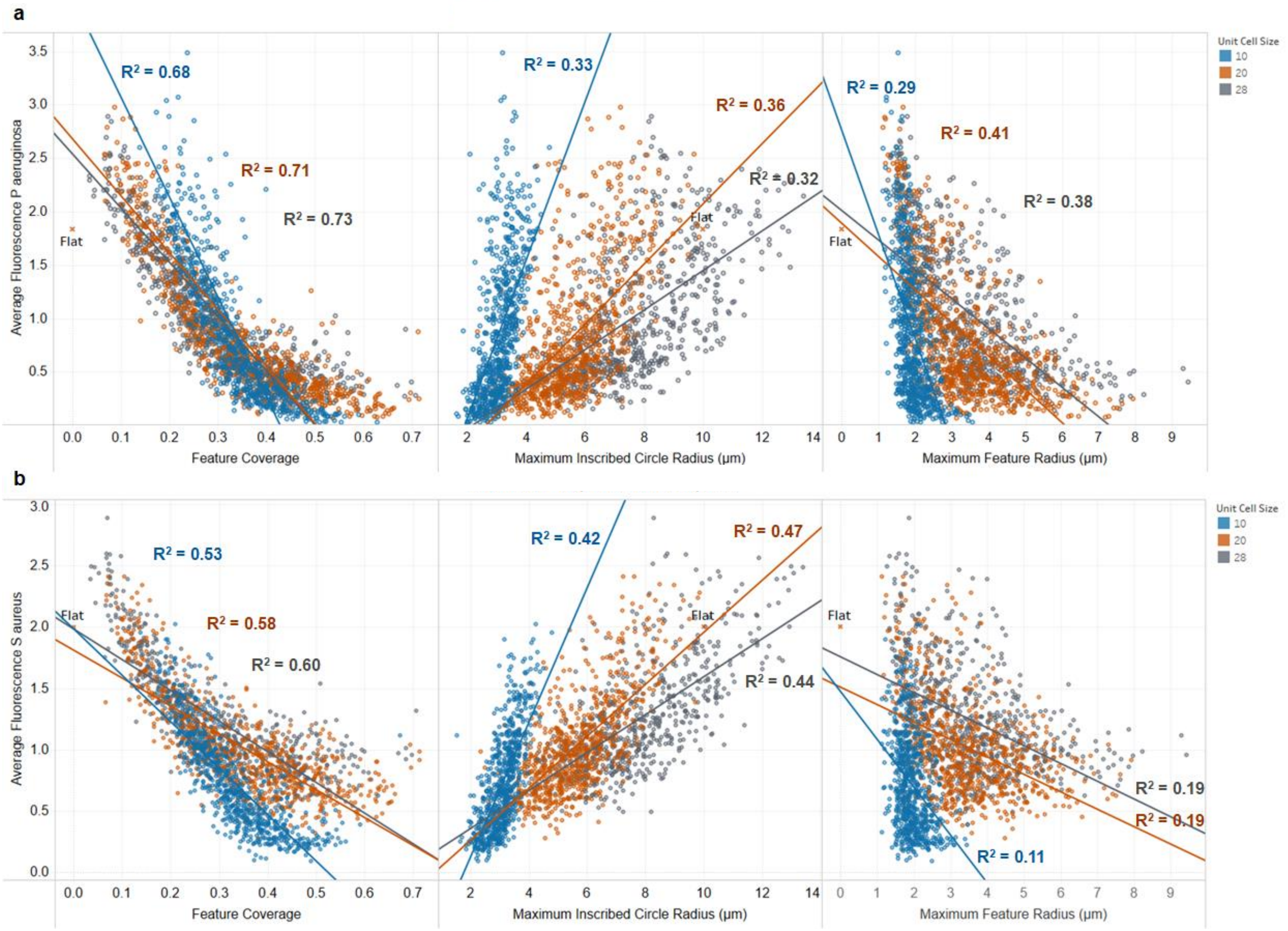


**Figure S4.** Topographical descriptors that show high correlation with bacterial attachment*:* (**a**) *P. aeruginosa* attachment; and (**b**) *S. aureus* attachment. The topographical descriptors found to be most important for both types of bacterial attachment are the feature coverage, the maximum size of inscribed circles radii, which relate to the space between the features and the radius of the largest feature in a topo unit cell.

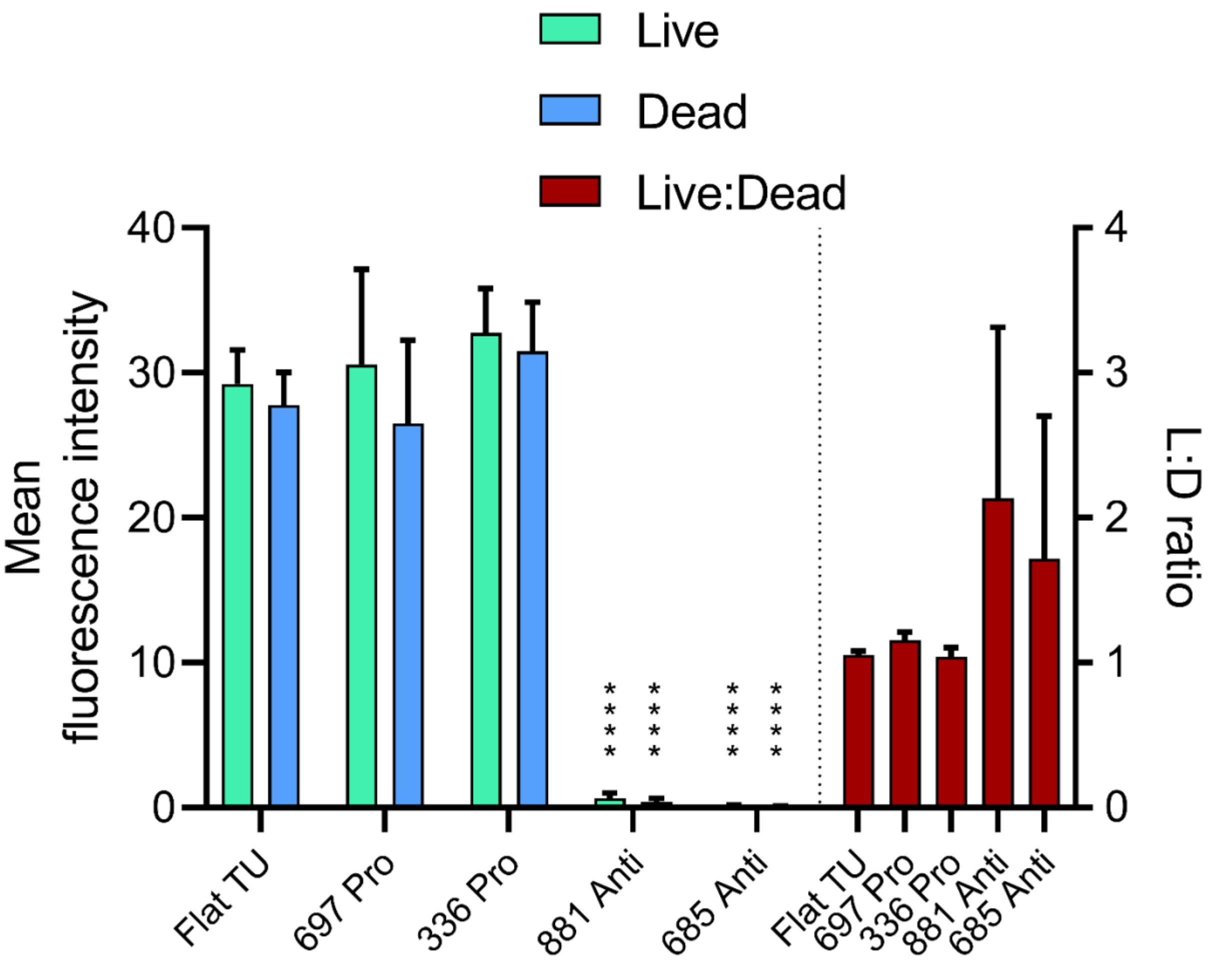


**Figure S5.** Mean fluorescence intensity of *P. aeruginosa* live/dead cells attached to flat, pro- (697 and 336) and anti-attachment (881 and 685) TUs after 4 h incubation in TSB + HS10% under static conditions. The ratios of live/dead cells attached to each TU are also shown (right Y-axis). Data shown are mean ±SD, n = 7. Statistical analysis was done using a two-way ANOVA with Dunnett's multiple comparisons test (* p<0.05; ** p<0.01; *** p<0.001; **** p<0.0001).

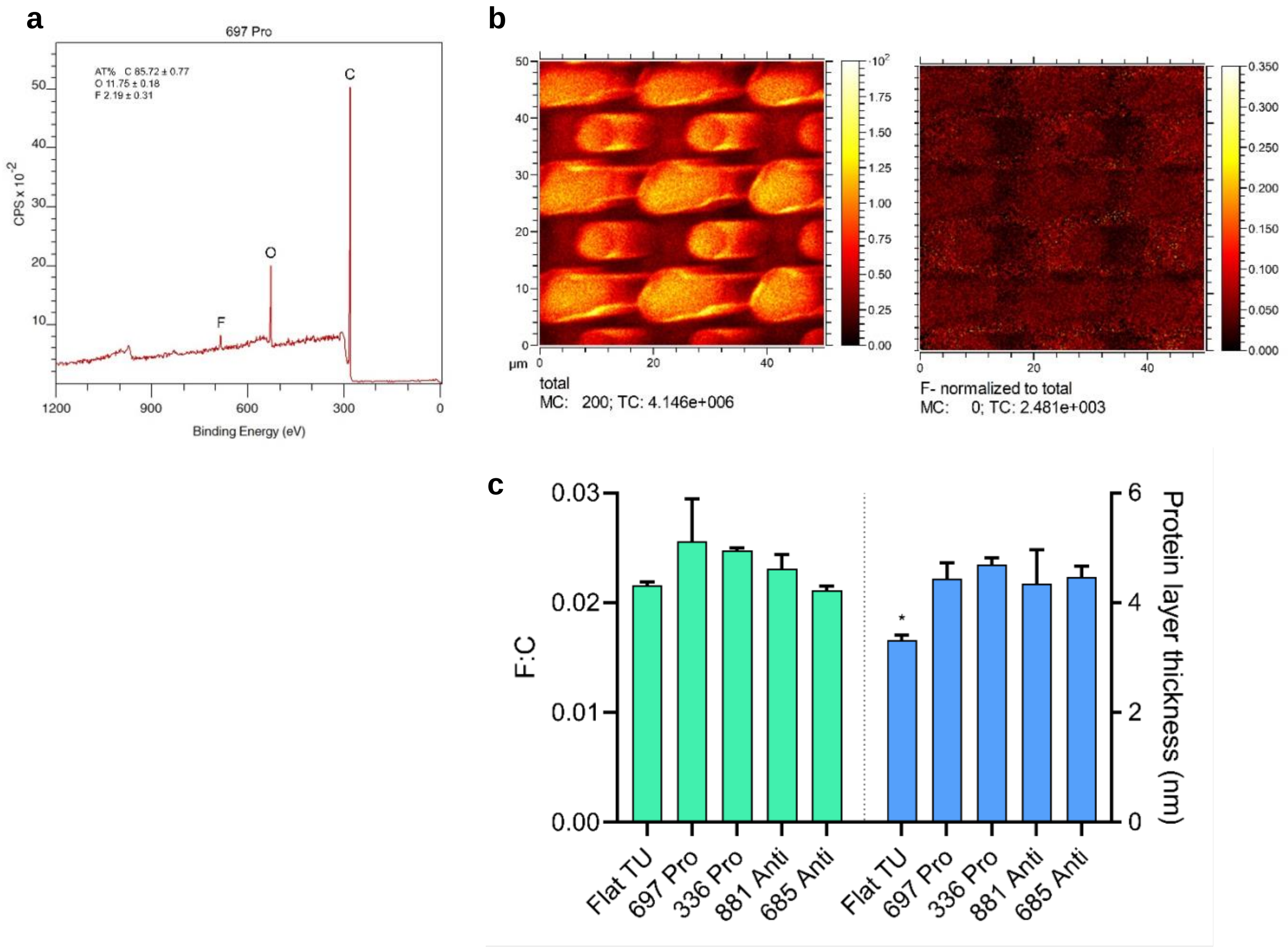


**Figure S6.** PS TopoChip surface chemistry analysis. (**a**) Representative XPS spectrum, obtained from a 100 × 100 μm area corresponding to a pro-attachment TU (697) in a plasma-treated chip, showing F impurity originated from PS TopoChip demoulding procedure. Atomic % for C, O and F elements are shown. (**b**) ToF-SIMS total negative and normalized F polarity secondary ion images obtained from a 50 x 50 μm area corresponding to TU (697) showing no differences in F content between topographical features of the same design. (**c**) XPS calculated F:C ratios from TUs with pro and anti-attachment properties against bacteria compared to flat surface control in PS TopoChip (green bars). Protein depth (nm) associated with pro and anti-attachment TUs and flat control after conditioning in TSB HS10% cell culture medium for 4 h (blue bars). Statistical differences between group means were determined by one-way ANOVA tests (*$p<0.05$).

**TopoChip surface chemistry**

As surface chemistry has a profound impact on bacterial attachment, it was essential to ensure that it was consistent across all the TUs. We therefore subjected the TopoChips to time-of-flight secondary ion mass spectrometry (ToF-SIMS) for molecular characterization with high lateral resolution together with X-ray photoelectron spectroscopy (XPS) for quantitative elemental analysis. Both methods detected fluorine containing impurities on the array surface, with XPS providing quantification for each TU on the array, e.g. topography 697 [F] = 2.2 ± 0.3 at% (**Fig S6a**). This could be assigned to residues from a monolayer of trichloro(1*H*,1*H*,2*H*,2*H*-perfluorooctyl)silane (FOTS) deposited on the OrmoStamp mould to facilitate moulding (Zhao et al., 2017). The distribution of F on the TU features, side walls and valleys was found to be constant using ToF-SIMS, within the limits of the technique imposed by the artefactual distortion of the features observed in **Fig. S6b**. Presenting a range of TUs where the F to C ratio was quantified by XPS, **Fig. S6c** illustrates that there was no statistically significant difference between the units (one-way ANOVA, $p>0.05$). These results indicate that the surfaces used in the screening have uniform chemistry and that the bacteria-material interactions observed are specifically dependent on surface topography.

Since TSB containing 10% serum (TSB HS10%) was used to simulate *in vivo* growth for some experiments, XPS analysis was carried out after incubation of TUs in uninoculated TSB HS10% medium for 4 h. No significant differences in the protein layer thicknesses were recorded between different TUs (**Fig. S6c**). Higher levels of nitrogen were detected on topographically defined surfaces compared to flat controls after TSB HS10% conditioning, corresponding with an increase in protein layer thickness (**Fig. S6c**). However, it is likely that the differences originate from the reduced sampling depth on the vertical feature sides that results in an over-estimation of protein layer thickness.

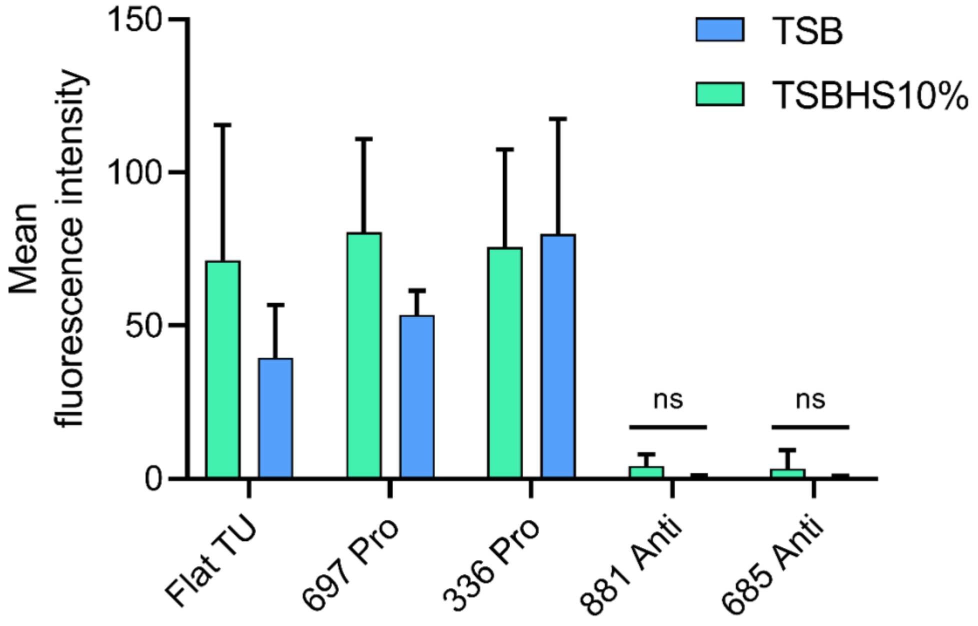


**Figure S7.** Comparative attachment of *P. aeruginosa* on flat, pro- (697 and 336) and anti-attachment (881 and 685) TUs in TSB or TSBHS 10% after 4 h incubation under static conditions. Data shown are mean ±SD, n ≥ 8. Statistical analysis was done using a two-way ANOVA with Dunnett's multiple comparisons test (ns $p$>0.05).

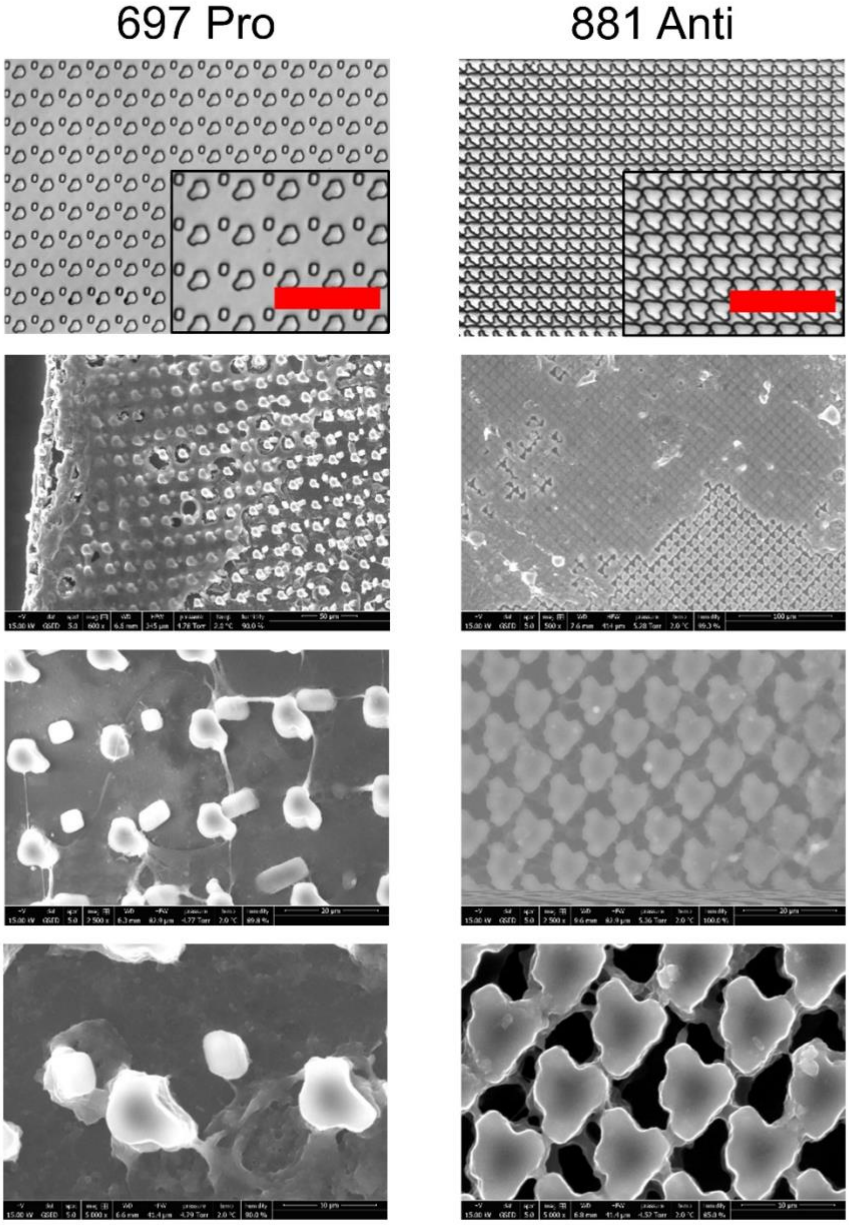


**Figure S8.** *Ex vivo* ESEM images of pro- (697) and anti-attachment (881) PU TUs removed from mice infected with *P. aeruginosa* for 4 days and imaged by ESEM. Scale bar in top row bright field images: 50 µm. ESEM image scales for 697 are 50, 20 and 10 µm; for 881, they are 100, 20 and 10 µm.
